# Multi-dimensional digital bioassay platform based on an air-sealed femtoliter reactor array device

**DOI:** 10.1101/2020.10.25.354381

**Authors:** Shingo Honda, Yoshihiro Minagawa, Hiroyuki Noji, Kazuhito V. Tabata

**Author notes:** Corresponding authors: Hiroyuki Noji and Kazuhito V. Tabata, **Email**. **Author Contributions** S.H., Y.M., H.N., and K.V.T. designed research; S.H. performed research and analyzed data; and S.H., H.N., and K.V.T. wrote the paper.

## Abstract

Single-molecule experiments have been helping us to get deeper inside biological phenomena by illuminating how individual molecules actually work. Digital bioassay, in which analyte molecules are individually confined in small compartments to be analyzed, is an emerging technology in single-molecule biology and applies to various biological entities (e.g., cells and virus particles). However, digital bioassay is not compatible with multi-conditional or multi-parametric assays, hindering understanding of analytes. This is because current digital bioassay lacks a repeatable solution-exchange system that keeps analytes inside compartments. To address this challenge, we developed a new digital bioassay platform with easy solution exchanges, called multi-dimensional (MD) digital bioassay, and tested its quantitativity and utility. We immobilized single analytes in arrayed femtoliter (10^−15^ L) reactors and sealed them with airflow. The solution in each reactor was stable and showed no cross-talk via solution leakage for more than 2 h, and over 30 rounds of perfect solution exchanges were successfully performed. To show the utility of our system, we investigated neuraminidase inhibitor (NAI) sensitivity on single influenza A virus (IAV) particles in a multi-conditional assay. We proved that IAV particles show a heterogeneous response to the NAI. Further, to demonstrate multi-parametric assays, we examined the sensitivity of individual IAV particles or model enzyme molecules to two different inhibitors. Our results support that MD digital bioassay is a versatile platform to unveil heterogeneities of biological entities in unprecedented resolution.

## Introduction

Although biochemical experiments in test tubes have long been the gold standard to analyze biological molecules, they only show us an averaged property of the population and do not allow us to access the individual molecules. In the field of single-molecule biology, various methods have been developed to detect signals from a single molecule (1).

Digital bioassay is one such method widely used in recent years (2–6). In this method, analytes (e.g., nucleic acids, enzymes, or other proteins) are diluted and partitioned into a large number of small compartments, which results in “0,” compartments containing no analytes, or “1,” compartments containing only a single analyte. Upon amplifying and detecting signals from these “digitalized” compartments, we can quantify analytes in a single-molecule resolution simply by counting the “1” compartments. Digital bioassay has enabled ultrasensitive detection of diverse analytes, including nucleic acids (called digital PCR [5, 7–11]), proteins (called digital ELISA [6, 12, 13]), and even influenza virus particles (14, 15). The most commonly used compartment in digital bioassay is a droplet emulsion (9, 10, 16, 17), typically a water droplet separated by a continuous oil phase. Arrayed reactors fabricated on a substrate (e.g., glass) (7, 8, 11–15, 18–26) are another type of frequently used compartment in which sample solutions are loaded into the reactors and sealed with oil (e.g., fluorocarbon oil).

Digital bioassay has also shown its potential to unveil heterogeneity of analytes. In single-molecule enzymology, it has revealed divergent activity states of certain hydrolytic enzymes such as β-galactosidase (18), β-glucuronidase (20), and alkaline phosphatase (14, 25, 26), by evaluating activities of the individual enzyme molecules partitioned in compartments.

However, current digital bioassays are limited by the fact that confined analytes cannot be analyzed more than twice and are incompatible with multi-dimensional (multi-conditional or multi-parametric) assays. This is because of the difficulty in exchanging solutions in the compartments while keeping the analytes inside. This limitation significantly hinders our understanding of analytes, as examining responses to varying conditions is one of the most standard approaches in biochemistry. For instance, we can obtain essential information such as enzyme-substrate affinity and catalytic efficiency from Michaelis-Menten parameters (27) determined by a series of enzyme activity measurements with changing substrate concentrations. Likewise, IC_50_ determined by a series of enzyme activity measurements with changing inhibitor concentrations is a standard indicator of inhibitor sensitivity (28). Although a determination of Michaelis-Menten parameters of β-galactosidase based on a single-molecule experiment has been reported (29), the parameters were not calculated based on the individual molecules. Another study measured the activity of single-molecule F_1_-ATPase with several different doses of substrate ATP (19). However, unlike in standard digital bioassay platforms, only a small number of analytes could be assayed in this system, which was insufficient for statistical analysis.

Although some methods are reported in droplet-based digital bioassay to inject other solutions into droplets or mix solutions in different droplets (30–33), it is still challenging to exchange solutions inside droplets without breaking them. In arrayed-reactor format, solution exchange can be performed by immobilizing an analyte on the surface of a reactor and washing out the solution inside after an assay. We once demonstrated a solution exchange in femtoliter (10^−15^ L) reactor array device (FRAD) fabricated on a glass substrate (26). We immobilized a model enzyme, alkaline phosphatase (ALP), in each reactor with adsorption and sealed the reactor with washable mineral oil, making solution exchange possible. However, repeated assays and solution exchanges with this approach resulted in reactors compromised with mineral oil, and the number of solution exchanges was fairly limited. Moreover, hydrophobic molecules (e.g., reporters, inhibitors, and effectors) inside the reactors tend to leak into hydrophobic sealing oil (21, 23, 24), including mineral oil, which leads to a decrease in the signal-to-noise ratio and hinders the quantitative assay. This is quite problematic because many of the fluorescent reporters used in digital bioassays are hydrophobic.

Here, we developed a new digital bioassay platform with easy solution exchanges, called multi-dimensional (MD) digital bioassay, by sealing reactors with airflow instead of oil, which also enabled leak-proof encapsulation. To test the quantitativity of our system, we confirmed that it meets three requirements: stability, independence, and exchangeability of solutions encapsulated in the reactors. Then we performed multi-conditional assays and multi-parametric assays to show its utility.

In a demonstration of a quantitative multi-conditional assay, we first conducted a single-particle analysis of the influenza A virus (IAV) immobilized in FRAD. Using hydrophobic fluorescent dye 4-methylumbelliferone (4-MU) as a reporter, we repeatedly measured the enzymatic activity of viral neuraminidase (NA) on individual IAV particles with the dose of NA inhibitor (NAI) gradually changed. We successfully determined the IC_50_s of an NAI for individual IAV particles. We also demonstrated a multi-parametric assay to estimate IC_50_s of two different NAIs on the same virus population. Finally, we applied our MD digital bioassay to single-molecule enzyme analysis and measured the IC_50_s of two inhibitors for a mixture of two ALPs from different origins. Based on the MD data obtained, we were able to distinguish one type of ALP from the other at single-molecule resolution. Thus, MD digital bioassay can offer a higher-resolution understanding of single biological entities such as virus particles and enzyme molecules than conventional digital bioassays.

## Results

### Digital bioassay based on air-sealed reactors and repeating solution exchanges

In MD digital bioassay, we used FRAD composed of more than 570,000 reactors (diameter; 3.3 μm, height; 0.4 μm, volume; 3.4 fL) fabricated on a 20 × 20 mm square surface with standard photolithography and dry etching (Supplementary Information, Materials and Methods, Device fabrication). We assembled a flow cell, as shown in Figure 1A, and performed sample loading, sealing with air, and solution exchanges, as shown in Figure 1B.

**Figure 1.**
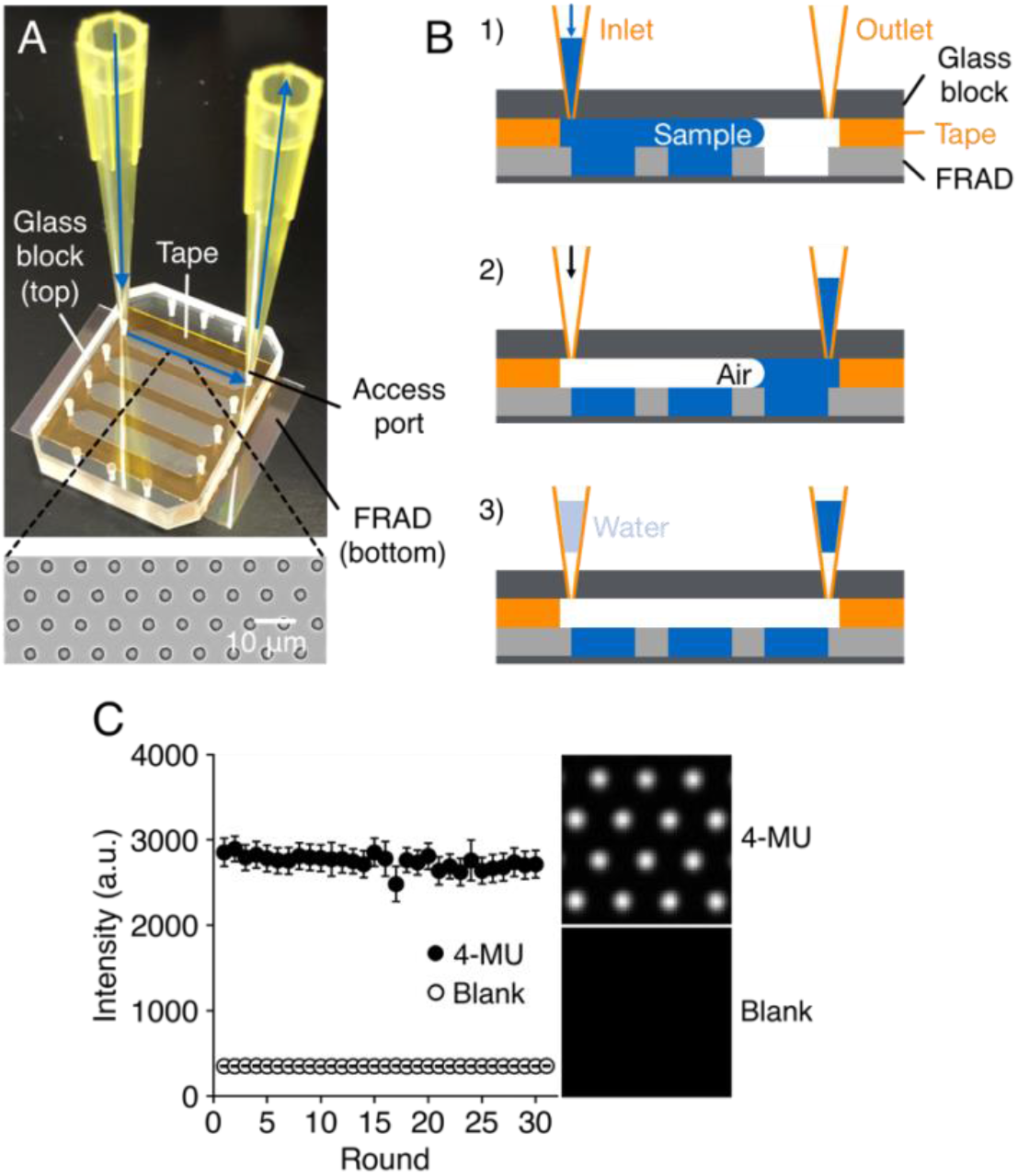
Sample encapsulation with airflow and solution exchanges. (A) Assembled flow cell. Solutions flow as shown by blue arrows. (B) Sample encapsulation procedure. 1) A sample is loaded into the flow channel. 2) Air is injected to push out excess sample and seal the reactors. 3) The flow channel is capped with water and sample at both ends to retain humidity and prevent evaporation. To exchange solutions, we replaced the solution inside with buffer or the next sample and repeated the sealing procedure. (C) Solution exchangeability of air-sealed reactors. The mean ± standard deviation (SD) of fluorescence intensity of >40,000 reactors are plotted for 30 rounds of solution exchanges. 4-MU (+): 1 mM 4-MU, 4-MU (-): blank buffer.

For the quantitative MD digital bioassay, the following requirements should be considered: solution stability (i.e., no evaporation from the reactors), (ii) solution independence (i.e., no cross-talk between the reactors), (iii) solution exchangeability (i.e., no fluorescent dye left inside the reactors after each round of solution exchange).

To investigate solution stability, we loaded fluorescent dye Alexa Fluor 647 into the reactors and sealed them with air to observe fluorescence from the reactors with a fluorescence microscope (Fig. S1). Without capping of the flow channel, we observed ring-shaped fluorescence within 2 h, indicating evaporation from the reactors (22). We quantified the degree of evaporation by plotting the fluorescence intensity at the center of each reactor over time. When we capped the flow channel, the intensity was around 100 % even after 2 h, similar to that in the oil-sealed flow channel as a positive control. Without capping, however, the intensity dropped down to 38 % after 2 h. This indicates that capping the flow channel prevent evaporation from the reactors and keep the solution stable for at least 2 h. We investigated possible cross-talk between the reactors via solution leakage by fluorescence recovery after photobleaching. We encapsulated Alexa Fluor 647 in the reactors with air and photobleached some of them with laser radiation. Fluorescence recovery was 0.1 % after 2 h, which was within the level of error among reactors (2.7 % relative standard deviation [RSD]) (Fig. S2), indicating that each reactor was independent and that no cross-talk between neighboring reactors occurred. Then, we tested whether we can exchange the solution inside the reactors completely. We measured the fluorescence intensity of the blank buffer encapsulated in the reactors (background). After washing it out, we encapsulated 4-MU and measured the fluorescence intensity (signal). We alternately measured the background and signal and then plotted the fluorescence intensity change (Fig. 1C). Rise in the background was 0.05 % after 30 rounds, which was within the level of error among reactors (2 % RSD), indicating that 4-MU was washed out thoroughly after each round for no less than 30 rounds of solution exchanges.

### Evaluation of enzymatic activity of single IAV particles using MD digital bioassay

To perform a quantitative analysis using our system, we evaluated the NA activity of IAV strain A/Puerto Rico/8/1934 (PR8) in a single-particle manner. We immobilized a single virus particle of PR8 in each reactor by adsorption (Fig. 2A). To ensure that only a single virus particle existed in each reactor, we diluted PR8 to the concentration where 0.02 particles on average were immobilized in a reactor. Although some reactors might contain more than two particles, we ignored them as they accounted for less than 0.02% of the total reactors. After washing out the unbound virus particles, we loaded 1 mM of a fluorogenic substrate, MUNANA, which is converted into fluorescent 4-MU by NA, and sealed the reactors with air. We measured the fluorescence intensity of each reactor over time and calculated the NA turnover of an individual virus particle from the conversion rate and the calibration curve (Fig. 2B, Fig. S3B, S4). Figure 2C shows the distribution of NA turnover. We defined the peak around 0 as background, which corresponded to the reactors with no virus particles. We calculated the mean and the standard deviation (SD) of the background peak by Gaussian fitting. We set a threshold (mean + 15SD, shown as an orange line in Fig. 2C) and excluded the reactors below it as background. We exchanged the solution and conducted three measurements on the same virus population to estimate measurement repeatability, i.e., intra-reactor error (Fig. 2D). The inter-reactor error was attributed to error from imaging system and variation in reactor volume and was estimated to be <5% RSD (Fig. S3C). Mean NA turnover was 670 turnovers/s/particle, and the variation was 33% to 39% RSD after subtraction of the measurement error consisting of intra-reactor error and inter-reactor error (Fig. S5, see also Supplementary Information, Materials and Methods, Data analysis, Section 4 for details).

**Figure 2.**
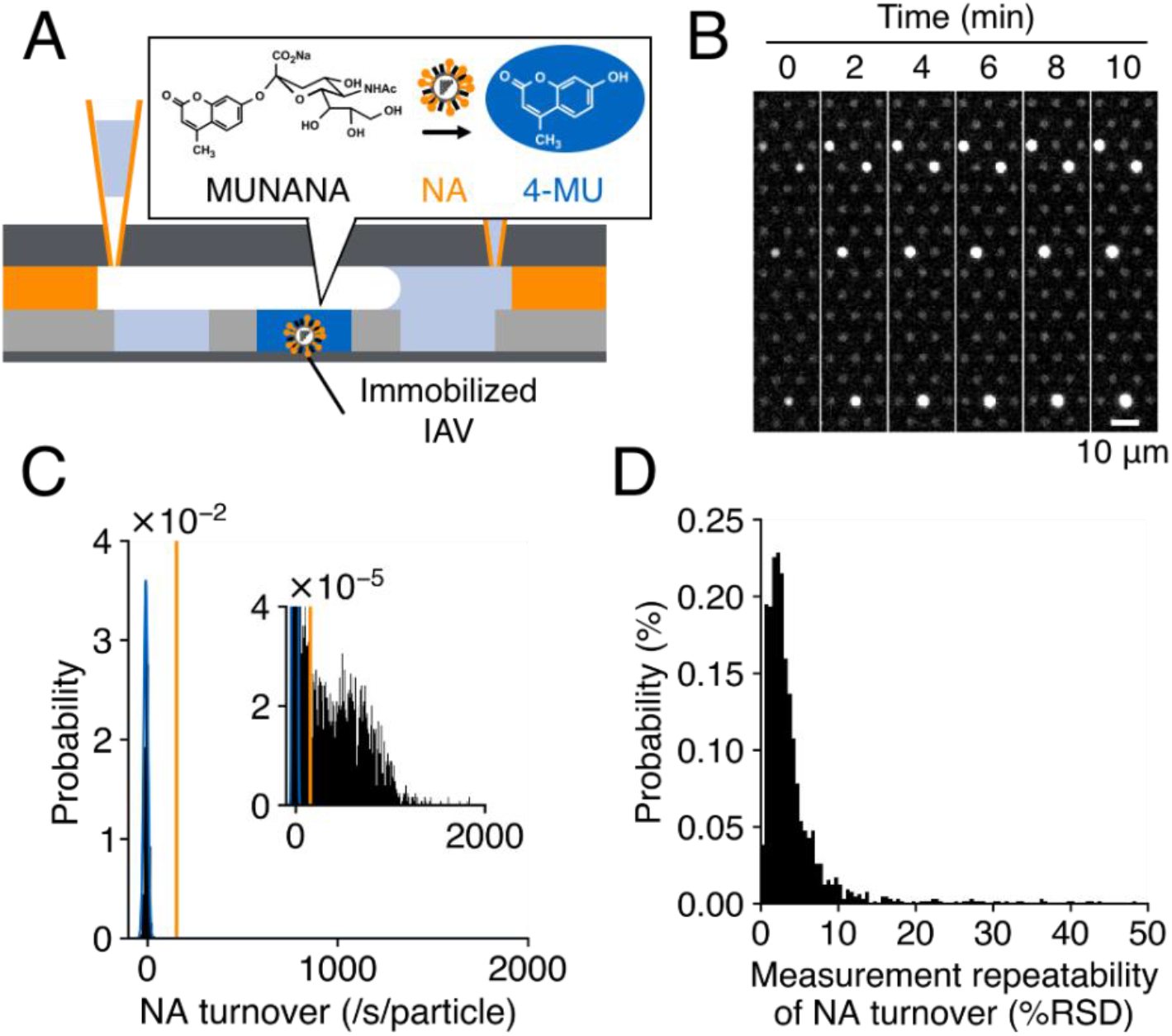
NA turnover analysis of individual IAV particles. (A) Schematic of the experiment. NAs on the surface of immobilized IAV convert MUNANA into 4-MU. We observed and quantified 4-MU accumulated in the reactor by a fluorescence microscope. (B) 4-MU fluorescent image of typical reactors at each time point. (C) Distribution of NA turnover. One result from the triplicate measurements is shown. Blue: Gaussian-fitted peak of reactors with no virus particles (background reactors). Orange: threshold to exclude background reactors (mean + 15SD). (D) Distribution of measurement repeatability of NA turnover measurements (*N* = 3 for each 1304 virus particles).

### Evaluation of NA inhibitor sensitivity of single IAV particles using MD digital bioassay

Based on the quantitative measurement of NA turnover, we evaluated the IC_50_ of NAI for individual virus particles of PR8 to demonstrate quantitative multi-conditional assay with solution exchanges. We repeatedly measured the NA turnover of individual virus particles of PR8 assayed, shown in Figure 2, gradually increasing the dose of NAI, oseltamivir (0, 1, 5, 20, and 500 nM). Then, we fitted the oseltamivir dose-dependent decrease in NA turnover with the Hill equation to calculate the IC_50_. We conducted 15 measurements in total, three times for five doses of oseltamivir, including three measurements in Figure 2, to estimate the measurement error. We observed dose-dependent decreases in NA turnover (Fig. 3A, 3B) and successfully calculated the IC_50_ for 1300 virus particles (Fig. 3B inset). The mean IC_50_ (3.9 nM) agreed well with that determined by conventional bulk scale bioassay on a microtiter plate (3.4 nM, shown as an orange line in Fig. 3B inset). Intra-reactor error reflected measurement repeatability and fitting error and gave the distribution shown in Figure 3C, while inter-reactor error was mostly canceled out in the process of fitting. We observed variation of IC_50_ that could not be explained by the intra-reactor error (*P*<0.001, one-way ANOVA) and estimated it to be 17% to 22% RSD after subtracting the intra-reactor error (Fig. S6 see also Supplementary Information, Materials and Methods, Data analysis, Section 5 for details). To test a potential correlation between the IC_50_ variation and the NA turnover variation, we conducted a correlation analysis. We found no correlation (Fig. 3D), suggesting the origins of these variations are distinct.

**Figure 3.**
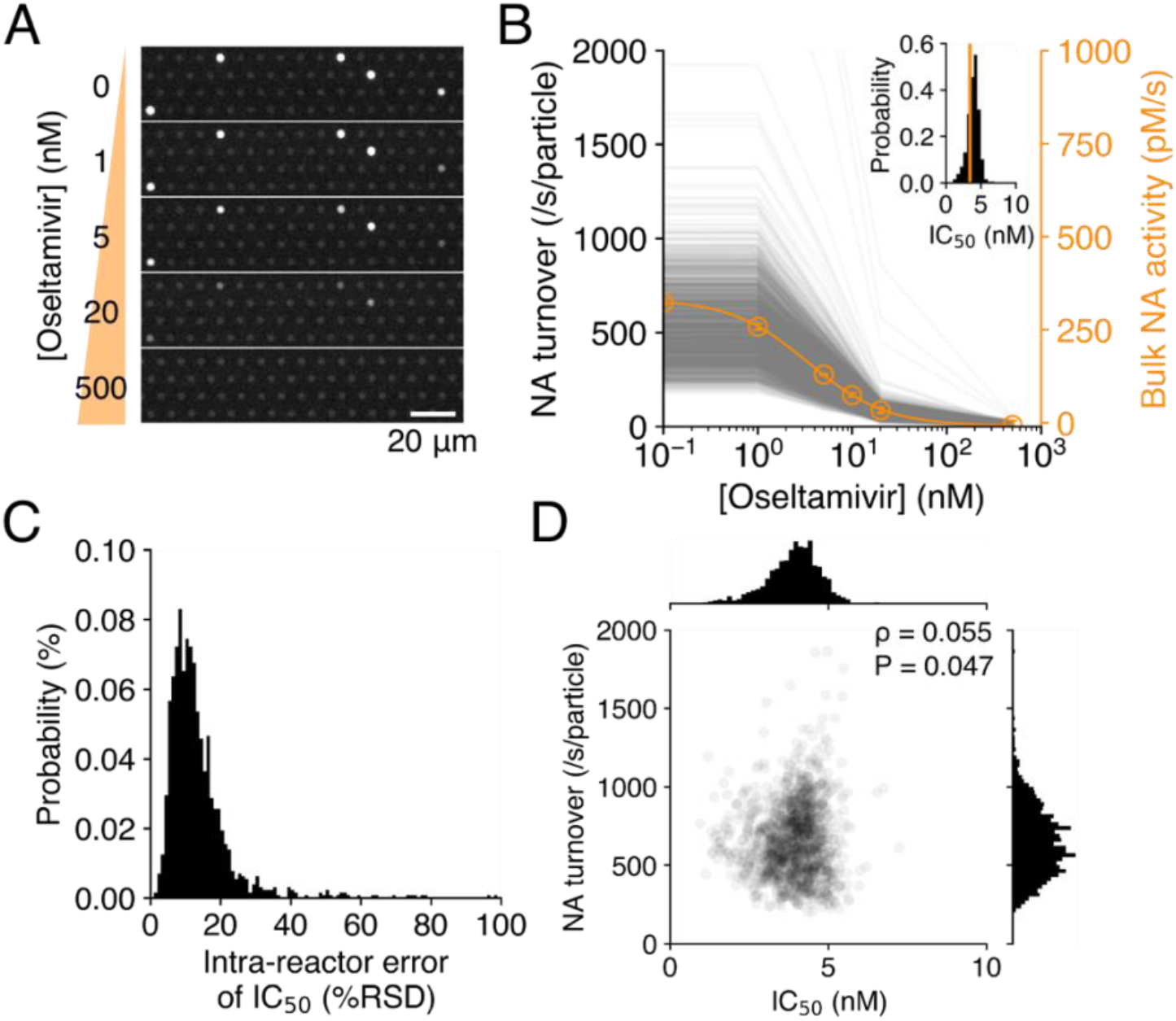
Oseltamivir sensitivity analysis of individual IAV particles. (A) 4-MU fluorescent image of typical reactors at different doses of oseltamivir (t = 10 min). (B) Oseltamivir dose-dependent NA turnover of 1300 virus particles. Each gray plot represents a single virus particle, and the mean of triplicate measurements in each reactor at each dose of oseltamivir is shown. Mean ± SD of NA turnover measured in a conventional bulk scale bioassay are plotted versus oseltamivir concentration and fitted with the Hill equation (orange curve). (inset) Distribution of IC_50_ of oseltamivir. Mean IC_50_ (3.4 nM) measured in a conventional bulk scale bioassay is indicated by the orange line. (C) Distribution of intra-reactor errors of IC_50_ measurements (1300 virus particles). (D) Correlation analysis between IC_50_ of oseltamivir and NA turnover. *ρ*: Spearman’s correlation coefficient.

### Multi-parametric assay of single virus particles and single enzyme molecules

Analysis with additional dimensions, such as multi-parametric assay, often allows us to gain deeper insight into analytes. To demonstrate multi-parametric assay with our system, we first evaluated the IC_50_s of oseltamivir and another NAI, zanamivir, on the same population of PR8. We immobilized PR8 in the reactors and measured NA turnover five times, gradually increasing the dose of oseltamivir (0, 1, 5, 20, and 500 nM). After clearing the reactors, we repeated the same procedure with zanamivir and calculated the IC_50_s of these two NAIs (Fig. 4A, Fig. S7). We observed no significant correlation between the IC_50_s of the two NAIs (Fig. 4B), suggesting that different factors determine their values.

**Figure 4.**
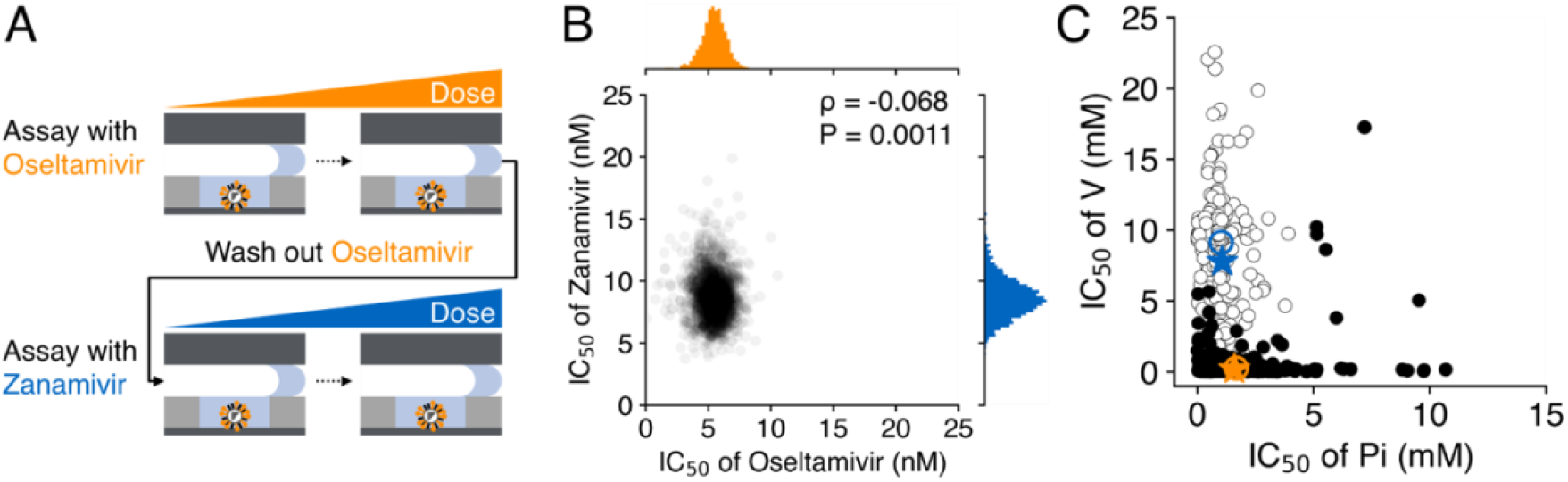
Multi-parametric analysis of single IAV particles and single enzyme molecules. (A) Schematic of the experiment with IAV particles. (B) Correlation analysis between IC_50_s of oseltamivir and zanamivir on single virus particles. *ρ*: Spearman’s correlation coefficient. (C) IC_50_s of Pi and V for single enzyme molecules of ALP mix (EcALP and CiALP). Blue and orange circles represent the centers of two fractions shown in white and black, respectively, resulting from the cluster analysis. Blue and orange stars represent the IC_50_s for EcALP and CiALP measured in conventional bulk scale bioassays.

To show the versatility of our system, we employed ALP as a model enzyme and performed another multi-parametric assay on individual ALP molecules. We focused on the fact that ALPs of different origins show distinct sensitivities to inhibitors (34, 35). We hypothesized that we could distinguish an ALP of one source from that of another in a mixture by cluster analysis based on the IC_50_s of inhibitors. To test this hypothesis, we conducted the following experiment using two ALPs (i.e., D101S, an active mutant of *Escherichia coli* ALP [EcALP] [23], and calf intestinal ALP [CiALP]) and two inhibitors (i.e., inhibitor phosphate [Pi] and vanadate [V]). We loaded the mixture of EcALP and CiALP into the flow channel, immobilized individual ALP molecules in the reactors, and washed out unbound ALPs. We used fluorogenic 4-methylumbelliferyl phosphate phosphatase substrate (4-MUP), which converts into 4-MU, to measure ALP turnover based on the conversion rate and calibration curve (Fig. S3A). We evaluated the IC_50_s of Pi and V for the ALP mix (Fig. S8) as we did for IAV. We employed two IC_50_s and the turnover rate as the parameters to conduct a cluster analysis (Fig. S9). As a result, we obtained two fractions, shown in Figure 4C. The IC_50_s of Pi and V at the centers were 1.0 mM and 9.1 mM in one fraction and 1.7 mM and 0.2 mM in the other. These were consistent with those of EcALP (1.1 mM, 7.8 mM) and CiALP (1.6 mM, 0.2 mM) measured in conventional bulk scale bioassays (Fig. S8), suggesting the two ALPs were correctly discriminated.

## Discussion

In this article, we established MD digital bioassay, a leak-proof digital bioassay platform with easy solution exchanges, by sealing reactors with airflow instead of oil commonly used in conventional digital bioassays. In a demonstration of multi-conditional assay, we quantitatively evaluated the IC_50_ of NAI for individual virus particles of IAV strain PR8 and revealed that PR8 shows heterogeneous sensitivity to NAI across the viral population. Further, we performed a multi-parametric assay on single virus particles of PR8 and single enzyme molecules of ALP.

We first confirmed the robustness of our system in terms of stability, independence, and exchangeability of solutions encapsulated in the reactors. The samples inside the reactors were stable and independent for at least 2 h and entirely exchanged no less than 30 times. Though we only performed assays within 10 min and under fewer than 15 conditions in this study, our system would also allow for longer times and higher-dimensional tests.

We then measured the NA turnover of individual virus particles of PR8 and estimated the measurement error, which consists of measurement repeatability and inter-reactor error. In our study, measurement repeatability was distributed around 2% RSD and was less than 10% RSD in 94% of the reactors (Fig. 2D). Compared under the same condition (50 μM 4-MU), the inter-reactor error in our system (<5.0% RSD, estimated from Fig. S3C) is superior to that in a previous study based on an oil-sealed femtoliter reactor array (7.1% RSD) (14). The variation in reactor volume should be less than 5.0% RSD under this condition, comparable to the best performance of droplet-based digital bioassays (5, 17). These errors are acceptable enough, at least for estimating the variation of a biological entity such as that addressed in this study (e.g., the NA turnover variation of individual IAV particles was approximately 33% to 39% RSD).

We calculated the NA turnover of single IAV particles as 670 turnovers/s/particle on average (25 °C). Given that an NA works as a tetramer and that a single IAV particle has ~50 tetramers (36, 37), we can estimate the NA turnover of a single IAV particle to be 1000 to 4000 turnovers/s/particle (37 °C) (38). This is reasonable because 670 turnovers/s/particle in 25 °C corresponds to 1000 to 1300 turnovers/s/particle in 37 °C considering the temperature change from 25 °C to 37 °C is reported to result in a 1.5- to 2.0-fold increase in NA activity (39). Our results demonstrate that MD digital bioassay can quantitatively evaluate NA turnover of individual virus particles. Because of its hydrophobicity and leakage into oil, the fluorescent dye 4-MU is challenging to use in digital bioassays even though it is a standard option to measure NA turnover with conventional bulk scale bioassays (40, 41). With oil-less sealing, our system demonstrated the first NA turnover measurement with a digital bioassay.

We estimated the variation of NA turnover to be 33% to 39% RSD. We previously freeze-dried a PR8 sample via the platinum-replica method and observed it with a transmission electron microscope to determine the diameter of single virus particles of PR8 (42). The diameter had a distribution (17% RSD) from which the variation in surface area can be estimated to be 34% RSD when we assume the particles are all spherical. This value is supported by another article (43). When the number of NA tetramers is proportional to the surface area and NA turnover is proportional to the number of NA tetramers, the variation in NA turnover also results in 34% RSD. Therefore, our results might reflect the virus size distribution.

Since NA is essential for budding of IAV, the NAI oseltamivir is most commonly used as an influenza medication (44). It is known that NAI sensitivity has certain heterogeneity among viral strains due to mutations around the active site (e.g., NAI-resistant mutants H274Y and N294S) (45). However, it remains unclear whether NAI sensitivity is heterogeneous among individual particles, despite its importance in understanding how resistant mutants emerge. This is because no methods are currently available to determine the IC_50_ for single virus particles.

In our study, the average IC_50_ of the NAI oseltamivir for single IAV particles was determined to be 3.9 nM (Fig. 3B), with a variation of 17% to 22% RSD (Fig. S6) that could not be explained by the measurement error, showing that PR8 is heterogeneous in oseltamivir sensitivity. The average agreed well with the 3.4 nM measured in a conventional bulk scale bioassay, suggesting that multi-conditional assay with our system is highly quantitative. The most feasible explanation for the IC_50_ variation is quasi-species. Because of the high mutation rate (e.g., 1.8 × 10^4^ substitutions per nucleotide per strand replicated for PR8 [46]) specific to RNA viruses, an IAV strain possesses various minor mutants called quasi-species (47). Moreover, some mutations around the NA active site are reported to affect sensitivity to NAI (45, 48), which might be demonstratable by sequencing a set of genomes of single IAV particles. We also investigated the IC_50_s of two different NAIs, oseltamivir and zanamivir, on single IAV particles as the first demonstration of a multi-parametric digital bioassay. We found that the IC_50_s of these two NAIs are almost independent (Fig. 4B). Although oseltamivir and zanamivir both target the NA active site and possess a similar structure, most of the reported resistance NA mutations affect IC_50_ of oseltamivir and zanamivir differently (49). This is partly because of structural differences between the two NAIs: pentyloxy group specific to oseltamivir (45) and guanidino group specific to zanamivir (50). Together with the “quasi-species” hypothesis, minor populations of mutants with this NAI-selective effect might contribute to the independence of two IC_50_s. While some articles have reported the characterization of NAI-resistant mutants in an IAV strain by next-generation sequencing (NGS) (47, 51, 52), NAI sensitivity itself has not been directly measured. Additionally, even NGS cannot offer single-particle resolution. Our system demonstrated the first quantitative analysis of NAI sensitivity in single-particle resolution. We could also measure the IC_50_s of multiple inhibitors at a time, which might be useful, for example, in detecting a mutant with selective resistance to a specific NAI.

Another feature of our system is its versatility. We applied it to a multi-parametric assay of single enzyme molecules and conducted a cluster analysis of a mixture of two sources of ALPs. ALP is expressed in various species as a homodimeric metalloenzyme and catalyzes the hydrolysis of phosphate monoesters under alkaline conditions. It is routinely used as a standard disease marker in clinical diagnosis (53). In particular, the expression pattern of ALP isozymes is reported to change with the development of certain types of cancers (54), underlining the importance of precise profiling of such isozymes.

We were able to distinguish one ALP from the other based on their different sensitivities to inhibitors (Fig. 4C). As mentioned above, our system is compatible with more than 30 rounds of solution exchanges, and higher dimensional analysis is also possible. Because a number of *in vivo* studies have reported diseases accompanied by a specific expression profile of enzymes (55), single-molecule clustering performed in this study might be an attractive option for clinical diagnostics. Recently, the identification of multiple enzymes in serum was reported based on single-molecule enzyme activity analysis and a set of fluorogenic substrates specific to each enzyme (24), leading to ultrasensitive disease biomarker detection. Another study showed that single-molecule activity analysis of ALP isozymes in serum could distinguish between patients with various health conditions (25). Although it remains to be determined how closely related enzymes can be distinguished with our system, it could detect even enzyme sub-populations undetectable with conventional bulk scale bioassays with a combination of these techniques. Further, our system would also be applicable to other biological entities (e.g., cells and extracellular vesicles) as long as a signal can be amplified from a single entity. Thus, MD digital bioassay is a versatile platform which can unveil heterogeneity of biological entities in unprecedented resolution.

## Materials and Methods

### Sample encapsulation with air

We used either assay buffer A (1 M DEA-HCl, 1 mM MgCl_2_, and 100 μM ZnSO_4_ [pH 9.25]) or assay buffer B (1 M DEA-HCl, 4 mM CaCl_2_ [pH 9.0]) in each experiment. We loaded an assay buffer for each experiment into a flow channel from the inlet and sonicated for a few seconds with an ultrasonic bath (US-2KS, SND) to remove air from the reactors. Then, we replaced the buffer with the sample and set a new pipette tip to the outlet. Air was injected at a rate of 1 μL/s with a micropipette to push excess sample out of the flow channel and seal the reactors. The ejected sample was used to cap the outlet to prevent evaporation from the reactors. The inlet was also capped with water.

### Enzyme activity analysis in FRAD

A stock of PR8 (1 × 10^11^ PFU/mL) was 1000-fold diluted with assay buffer B. Stocks of EcALP and CiALP were 4.0 × 10^5^-fold and 2.3 × 10^5^-fold diluted, respectively, with assay buffer A. Then, we mixed 2 μL each of the two diluted ALPs and added 16 μL of assay buffer A. We loaded the diluted PR8 or ALP mix into a flow channel and incubated it at 20 °C to 25 °C for 30 min to immobilize it with adsorption. For the ALP mix, we sealed the reactor with air before incubation. We washed the flow channel 10 times with the buffer and mounted the flow cell on the microscope stage. We then conducted enzyme activity analyses or inhibitor sensitivity analyses. All analyses were conducted at 25 °C.

In the NA activity analysis of PR8 (Fig. 2, Fig. S4, S5) and the ALP activity analysis (Fig. 4C, Fig. S8), we loaded 1 mM MUNANA in assay buffer B containing 10 μM Alexa Fluor 647 and 1 mM 4-MUP in assay buffer A containing 10 μM Alexa Fluor 647, respectively, into the flow channel. We sealed the reactor with air and recorded fluorescent images of 4-MU and Alexa Fluor 647 every 2 min for 10 min. We washed the flow channel with the buffer, loaded another substrate, and repeated the measurement.

### Inhibitor sensitivity analysis in FRAD

We conducted an oseltamivir sensitivity analysis of PR8 (Fig. 3, Fig. S6, S7) on the same virus population in which we analyzed NA activity (Fig. 2). We loaded 1 mM MUNANA in assay buffer B containing 10 μM Alexa Fluor 647 and NAI into the flow channel. After 1 h of preincubation to ensure that NAs were saturated with oseltamivir, we sealed the reactors with air and recorded fluorescent images of 4-MU and Alexa Fluor 647 every 2 min for 10 min. We loaded another substrate and oseltamivir and repeated the measurement, gradually increasing the dose of oseltamivir (0, 1, 5, 20, and 500 nM). In the multi-parametric assay with two NAIs (Fig. 4B, Fig. S7), we conducted an oseltamivir sensitivity analysis and washed out oseltamivir bound to NAs as follows: 1) We washed the flow channel twice with the buffer and incubated it at 37 °C for 1 h using a stage heater (TIZB-B13F, Tokai Hit) mounted on a microscope (Eclipse Ti2-E, Nikon); 2) we repeated step 1 twice and washed the flow channel with the buffer. After washing out oseltamivir, we conducted a zanamivir sensitivity analysis. Then, we washed out zanamivir in the same manner and analyzed the NA activity of PR8 to identify reactors that had retained virus particles throughout the analyses.

The inhibitor sensitivity analysis of ALPs was performed in a similar manner (Fig. 4C, Fig. S8). We loaded 1 mM 4-MUP in assay buffer A containing 10 μM Alexa Fluor 647 and Pi into the flow channel. We sealed the reactors with air and recorded fluorescent images of 4-MU and Alexa Fluor 647 every 2 min for 10 min. We loaded another substrate and Pi and repeated the measurement, gradually increasing the dose of Pi (0, 0.01, 0.1, 1, 10, 100 mM). After washing the flow channel with the buffer, we repeated the analysis with another inhibitor, V. Then we washed the flow channel with buffer and analyzed the ALP activity to identify reactors that had retained ALP molecules throughout the analyses.

## Acknowledgments

We thank Dr. H. Yaginuma and R. Kobayashi for helpful discussions; K. Ohtake for virus sample preparation; and all members of the H.N. laboratory for valuable comments. This work was supported by Grant-in-Aid for Scientific Research (S) (JP19H05624) to H.N., JST CREST, Japan (JPMJCR19S4) to H.N., the Japan Science and Technology Agency for Core Research for Evolutional Science and Technology (CREST) (JPMJCR18S6) to K. V. T., Program for Creating STart-ups from Advanced Research and Technology (START) (JPMJST1816) to K. V. T., and the MEXT Scientific Research on Innovative Areas, ‘Chemistry for multimolecular crowding biosystem’ (17H06355) to K. V. T..

## Supplementary Information

## Materials and Methods

### Materials

Diethanolamine (DEA), CaCl_2_-2H_2_O, 6N HCl, MgCl_2_-6H_2_O, TWEEN20, and K_2_HPO_4_ were purchased from FUJIFILM Wako Pure Chemical. ZnSO_4_-7H_2_O, 4-methylumbelliferyl phosphate phosphatase substrate (4-MUP), 2’-(4-methylumbelliferyl)-α-D-*N*-acetylneuraminic acid sodium salt hydrate (MUNANA), AZ P4903, and AZ 300 MIF Developer were purchased from Merck. Fluorinert™ FC-40 was purchased from 3M. Fomblin Y LVAC 25/6 was purchased from Solvay. CYTOP (CTL-809M) was purchased from AGC. Oseltamivir acid and zanamivir were purchased from ChemScene. Na_3_VO_4_ was purchased from Santa Cruz Biotechnology. PURExpress In Vitro Protein Synthesis Kit and PURExpress Disulfide Bond Enhancer (PDBE) were purchased from New England Biolabs. Recombinant RNase Inhibitor (RRI) was purchased from Takara Bio. Alexa Fluor 647 dye and 4-methylumbelliferone were purchased from Thermo Fisher Scientific and TCI, respectively. Calf intestinal alkaline phosphatase (CiALP; Alkaline Phosphatase recombinant, highly active) was purchased from Roche. Influenza A virus A/Puerto Rico/8/1934 (H1N1) was prepared as described previously (1). The plasmid coding D101S mutant of alkaline phosphatase (ALP) from *Escherichia coli* (EcALP) was prepared as described previously (2).

### Device fabrication

Femtoliter reactor array device (FRAD) was fabricated as described below. We dipped a 24 mm × 32 mm coverslip (Matsunami) into 8N KOH and sonicated it for 15 min in an ultrasonic bath (UT-206, Sharp). It was rinsed with deionized water and dried with blown air. Fluoropolymer CYTOP was spin-coated on the coverslip at 4000 rpm for 30 s and baked at 90 °C for 10 min followed by 180 °C for 30 min. Positive photoresist AZ P4903 was spin-coated on the CYTOP layer at 7500 rpm for 30 s and baked at 55 °C for 3 min followed by 110 °C for 5 min. Using a mask aligner (B100 IT, Nanometric Technology), the photoresist layer was exposed to UV light through a chrome-coated photomask patterned with arrayed pores (3 μm in diameter) with a pitch of 9 μm. The exposed photoresist was developed in AZ 300 MIF developer for 5 min, rinsed with deionized water, dried with blown air, and used as a mask for the following dry etching procedure. The masked CYTOP layer was etched with O_2_ plasma generated by a reactive ion etching system (RIE-10NR, Samco), resulting in a femtoliter reactor array. We sonicated the dry-etched device twice in acetone for 10 min and once in 2-propanol for 5 min, rinsed it with deionized water, and dried it with blown air. We measured the size of the reactors with a 3D laser scanning microscope (VK-X200, Keyence). We then assembled a flow cell by sticking a glass block on FRAD with double-sided tape (7602 #25, Teraoka) cut into the shape of flow channels. Before the assembly, CYTOP was spin-coated on the glass block at 1000 rpm for 30 s and baked at 180 °C for 1 h.

### Imaging

All images except those in the fluorescence recovery after photobleaching (FRAP) experiment (Fig. S2) were acquired with an inverted fluorescence microscope (Eclipse Ti2-E, Nikon) with an LED light source (X-Cite TURBO, Excelitas Technologies), appropriate filter sets (λ_ex_ = 360 nm and λ_em_ = 460 nm for 4-MU; λ_ex_ = 638 nm and λ_em_ = 670 nm for Alexa Fluor 647; Nikon), and an sCMOS detector (Zyla, Andor). In the FRAP experiment, we used a confocal laser microscope system (A1R, Nikon) with an appropriate laser and filter set for Alexa Fluor 647 and photomultiplier tubes. We used a 20× objective lens in all experiments except in the validation of solution stability (Fig. S1) and the FRAP experiment (Fig. S2), in which 40× and 60× objective lenses, respectively, were used.

### Validation of solution stability and independence

We encapsulated 50 μM Alexa Fluor 647 dye in assay buffer A in FRAD with air. In the validation of solution stability, we also prepared a non-capped flow channel as the negative control and a flow channel sealed with perfluorocarbon oil FC-40 as the positive control. From 1 min after sealing, we recorded fluorescent images of Alexa Fluor 647 every 15 min for 2 h. In the validation of solution independence, we photobleached 10 reactors for a few seconds at λ = 640 nm with a confocal laser microscope and recorded fluorescent images of Alexa Fluor 647 every 5 min for 2 h.

### Validation of solution exchangeability

We encapsulated a “blank” sample (10 μM Alexa Fluor 647 in assay buffer B) in FRAD with air and recorded a fluorescent image of 4-MU. Then, we washed the flow channel with assay buffer B. We encapsulated a “signal” sample (1 mM 4-MU in assay buffer B with 10 μM Alexa Fluor 647), recorded a fluorescent image of 4-MU, and washed the flow channel. We repeated this for 30 rounds.

### Cell-free protein synthesis of D101S, an active mutant of *E. coli* ALP

We synthesized EcALP using PURExpress In Vitro Protein Synthesis Kit according to the manufacturer’s instructions with some modifications. The reaction mixture (6 μL of solution A, 4.5 μL of solution B, 0.6 μL of PDBE1, 0.6 μL of PDBE2, 0.3 μL of RRI, 1.5 μL of 1 mM ZnSO_4_, 20 ng of template DNA, and ultra-pure water up to 15 μL) was incubated at 27 °C for 4 h. The resultant mixture was 1000-fold diluted with assay buffer A and used as a stock solution.

### Enzyme activity analysis and inhibitor sensitivity analysis on a microtiter plate

We performed all analyses at 25 °C on a 384-well black plate (781906, Greiner Bio-One). A stock of PR8 (1 × 10^11^ PFU/mL) was 100-fold diluted with 1 mM MUNANA in assay buffer B containing 10 μM Alexa Fluor 647 and NAI. The fluorescence intensity of 4-MU in 15 μL of the diluted PR8 was measured in a plate reader (Spectra Max iD3, Molecular Devices) every 5 min for 5 h (λ_ex_ = 360 nm, λ_em_ = 448 nm). Stocks of EcALP and CiALP were 2.8 × 10^4^-fold and 4.4 × 10^7^-fold diluted, respectively, with 1 mM 4-MUP in assay buffer A containing 10 μM Alexa Fluor 647, 0.02% (v/v) TWEEN20, and an inhibitor (Pi or V). Note that 4-MUP was added after 30 min of preincubation of the diluted ALP with an inhibitor at 20 °C to 25 °C. The fluorescence intensity of 4-MU in 40 μL of the diluted ALP was measured in a plate reader every 1 min for 15 min (λ_ex_ = 360 nm, λ_em_ = 448 nm). To generate a calibration curve for 4-MU quantification, we measured the fluorescence intensity of 4-MU diluted with the same volume of each assay buffer used in the analysis.

### Data analysis

#### 1) Calculation of fluorescence intensity of a dye in a reactor

We calculated the fluorescence intensity of 4-MU and Alexa Fluor 647 in each reactor recorded in each experiment using a custom ImageJ macro. Briefly, we applied a Gaussian filter to the fluorescent images of Alexa Fluor 647 and defined the top of each peak corresponding to each reactor as the center of the reactor. We drew a 10 pixels × 10 pixels square around this center to surround the reactor. The fluorescence intensity of 4-MU or Alexa Fluor 647 averaged over the surrounded area was defined as the mean fluorescence intensity of the dye in the reactor. Note that, in the validation of solution stability, the fluorescence intensity at the central pixel was defined as the fluorescence intensity of the dye in the reactor.

#### 2) Validation of solution stability, independence, and exchangeability

In the validation of solution stability (Fig. S1), the fluorescence intensity of Alexa Fluor 647 in each reactor at each time point was normalized to that at t = 0 min. The mean ± standard deviation (SD) of the normalized fluorescence intensity of Alexa Fluor 647 in the reactors was plotted versus time.

In the validation of solution independence (Fig. S2), the mean ± SD of the fluorescence intensity of Alexa Fluor 647 of the reactors was plotted versus time before and after photobleaching.

In the validation of solution exchangeability (Fig. 1C), the distribution of the fluorescence intensity of 4-MU in the reactors was fitted by a Gaussian function to calculate the mean ± SD. The mean ± SD was plotted versus the rounds of solution exchanges.

#### 3) Enzyme activity analysis in FRAD

In the NA and ALP turnover measurements, we excluded improperly sealed reactors, typically fused to large droplets formed outside the reactors, from the subsequent analyses as follows: The distribution of the fluorescence intensity of Alexa Fluor 647 in the reactors was fitted with a Gaussian function to calculate the mean ± SD. Since improperly sealed reactors show higher fluorescence intensity than properly sealed ones, we set a threshold (mean + 3SD) and excluded the reactors above this.

We converted the fluorescence intensity of 4-MU to the number of 4-MU molecules in a reactor using the calibration curve of 4-MU (Fig. S3A and S3B) and the reactor volume (3.4 fL). The numbers of 4-MU molecules at t = 0, 2, and 4 min were fitted with the least-square method to calculate the slope, i.e., the turnover of a single entity (IAV particle or ALP molecule). We plotted histograms for the distributions of NA and ALP turnover and defined the peak around 0 as background, which corresponds to the reactors without entities. We calculated the mean ± SD of the background peak by Gaussian fitting. We set a threshold (NA, mean + 15SD; ALP, mean + 3SD) in all the turnover measurements without an inhibitor and excluded the reactors below it as background, i.e., without entities. For the subsequent analyses, we used the reactors fulfilling the threshold in all the measurements without an inhibitor (1304 reactors in Figs. 2, 3, S5, and S6; 2286 reactors in Figs. 4B and S7; 2200 reactors in Figs. 4C and S8). In each reactor and at each dose of NAI, mean, SD, and RSD of NA turnover were calculated from triplicate measurements.

#### 4) Measurement error estimation of NA turnover

We estimated measurement error and intrinsic variation of NA turnover among 1304 IAV particles with one-way ANOVA (Figs. 2D, S5). To prepare sufficient datasets for ANOVA, we employed bootstrapping. We re-sampled the NA turnover measured with *i* (nM) oseltamivir (*i* = 0, 1, 5, 20, 500) in the *j*th reactor (*j* = 1, 2, 3, …, 1304) 100 times for each *i* and *j*. The detailed procedure is as follows: Based on the mean ± SD of NA turnover determined for a set of *i* and *j* from the triplicate measurements, we generated a random number following a normal distribution N (mean, SD^2^). We repeated this 100 times and named the *k*th random number generated as “v*_i, j, k_*” (*k* = 1, 2, 3, …, 100). v*_i, j, k_* (*k* = 1, 2, 3, …, 100) was employed as a virtual NA turnover dataset for each *i* and *j*.

We used v_0_*_, j, k_* for the measurement error estimation of NA turnover. The variance of v_0, j, k_ consists of intra-reactor error (among v_0,*j*, 1_, v_0, *j*, 2_, v_0, *j*, 3_, … v_0, *j*, 100_) and inter-reactor variation (among v_0, 1, *k*_, v_0, 2, *k*_, v_0, 3_, *k*, …, v_0, 1304, *k*_). The intra-reactor error is equivalent to measurement repeatability, and the inter-reactor variation is derived from intrinsic variation and inter-reactor error among 1304 virus particles. The inter-reactor error is attributed to error from imaging system and variation in reactor volume.

Although we first tried to separate the inter-reactor variation from the intra-reactor error with one-way ANOVA, we could not because the measurement repeatability differed among the reactors (Fig. 2D), which meant the intra-reactor error (SD) of NA turnover in each reactor was not homogeneous. Therefore, we “sliced” 1304 virus particles into subpopulations by the SD of NA turnover in each reactor at an SD increment of 4 turnovers/s/particle. Then, one-way ANOVA was applied to each subpopulation (Fig. S5), assuming that the SD of NA turnover in the subpopulation in each reactor was homogeneous. We estimated the inter-reactor variation (orange circles in Fig. S5) separate from the intra-reactor error (blue circles in Fig. S5). We regarded the estimated inter-reactor variation in subpopulations with more than 10% of the original virus particle population (≥131, closed orange circles in Fig. S5) as sufficiently reliable (33% to 39% RSD). The inter-reactor error depends on the fluorescence intensity of 4-MU in each reactor (Fig. S3C). As mentioned above, NA turnover was determined by fitting fluorescence intensity of 4-MU in a reactor at t = 0 to 4 min. Given that the fluorescence intensity at t = 4 min (700 ± 130) was the highest and therefore mostly controlled the fitting, the inter-reactor error can be estimated to be 5% RSD at most (Fig. S3C). Even after subtraction of this inter-reactor error from the inter-reactor variation, the intrinsic variation among the virus particles was 33% to 39% RSD, which suggested the inter-reactor error had little impact on the result.

#### 5) NAI sensitivity analysis in FRAD and the measurement error estimation

We determined IC_50_s of NAI on individual IAV particles and estimated the measurement error and the intrinsic variation among 1304 IAV particles with one-way ANOVA (Figs. 3B inset, 3C, S6). For this purpose, we used the virtual NA turnover dataset v_i, j, k_, where *i, j*, and *k* stand for the dose of oseltamivir (*i* = 0, 1, 5, 20, 500), reactor number (*j* = 1, 2, 3, …, 1304), and re-sampling number (*k* = 1, 2, 3, …, 100), respectively. We fitted ([v_0, *j, k*_], [v_1, *j, k*_], [v_5, *j, k*_], [v_20, *j, k*_], [v_500, *j, k*_]) with the Hill equation and obtained an IC_50_ value for each *j* and *k*, which we named IC_50, *j, k*_. After excluding four reactors that we failed to fit, we calculated the mean (Fig. 3B inset), SD, and RSD (Fig. 3C) of the IC_50_ values (IC_50, *j*_, 1, IC_50, *j*, 2_, IC_50, *j*, 3_, …, IC_50, *j*, 100_) for each *j*. The variance of IC_50, *j, k*_ consists of intra-reactor error (among IC_50, *j*, 1_, IC_50, *j*, 2_, IC_50, *j*, 3_, … IC_50, *j*, 100_) and inter-reactor variation (among IC_50, 1, *k*_, IC_50, 2, *k*_, IC_50, 3, *k*_, …, IC_50, 1300, *k*_). The intra-reactor error is equivalent to measurement repeatability, including fitting error, and the inter-reactor variation directly reflects the intrinsic variation of IC_50_ among the 1300 virus particles, as the inter-reactor error is mostly canceled out in the process of fitting. As measurement repeatability differed among the reactors (Fig. 3C), we “sliced” the 1300 virus particles into subpopulations by the SD of IC_50_ in each reactor at an SD increment of 0.1 nM, as we did with NA turnover in the previous experiment. Then, one-way ANOVA was applied to each subpopulation (Fig. S6), assuming that the SD of IC_50_ in the subpopulation in each reactor was homogeneous. We estimated the intrinsic variation (orange circles in Fig. S6) separate from the intra-reactor error (blue circles in Fig. S6). We regarded the estimated intrinsic variation in subpopulations with more than 10% of the original virus particle population (≥131, closed orange circles in Fig. S6) as sufficiently reliable (17% to 22% RSD). The intrinsic variation of IC_50_ in all these subpopulations were significant and could not be explained by the measurement error (*P*<0.001, one-way ANOVA).

#### 6) ALP inhibitor sensitivity analysis in FRAD

We determined IC_50_s of ALP inhibitors (Pi and V) for individual ALP molecules by fitting the ALP turnovers measured at each dose of an inhibitor (0, 0.01, 0.1, 1, 10, and 100 mM) in each reactor (2200 reactors) with the Hill equation. Variation of calculated IC_50_s for individual ALP molecules was higher than that calculated for individual IAV particles. Moreover, we failed to fit a greater number of reactors than we did in the case of IAV particles. This was likely due to the stochastic feature of a single-molecule experiment. A single-molecule ALP turnover measurement is usually accompanied by the high variability caused by stochastic fluctuation in the enzyme structure, while in the NA turnover measurements on a single IAV particle, an IAV particle has approximately 50 NA tetramers; NA turnover was averaged over the particle, which reduced the variation. To mitigate the impact of the IC_50_ variability on the subsequent cluster analysis, we set a threshold (the coefficient of determination for the fitting, R^2^ > 0.9 and IC_50_ < 25 mM) and excluded the reactors that did not meet this threshold. Note that the IC_50_ variability was considered to be mainly due to the fitting error caused by the random variation in ALP turnover measurements, which means it is unlikely that the thresholding caused certain biases in the ALP population.

#### 7) Enzyme activity analysis and inhibitor sensitivity analysis on a microtiter plate

We calculated the 4-MU concentrations of the samples at each time point from the 4-MU fluorescence intensity and the calibration curves (Fig. S3A, B). Then, we fitted the 4-MU concentrations at 1 to 5 h (NA) or at 0 to 15 min (ALP) with the least-square method to determine the enzyme activity. Enzyme activity at each dose of inhibitor was fitted with the Hill equation to determine IC_50_ (Figs. 3B, S7, S8).

#### 8) Cluster analysis of ALP mix

We normalized the ALP turnover determined without an inhibitor and the IC_50_s of Pi and V in each reactor and used them as input parameters for cluster analysis. With the hierarchical Ward’s method (3), we separated the population of ALP mix into two fractions (Figs. 4C, S9).

---

**Fig. S1.**
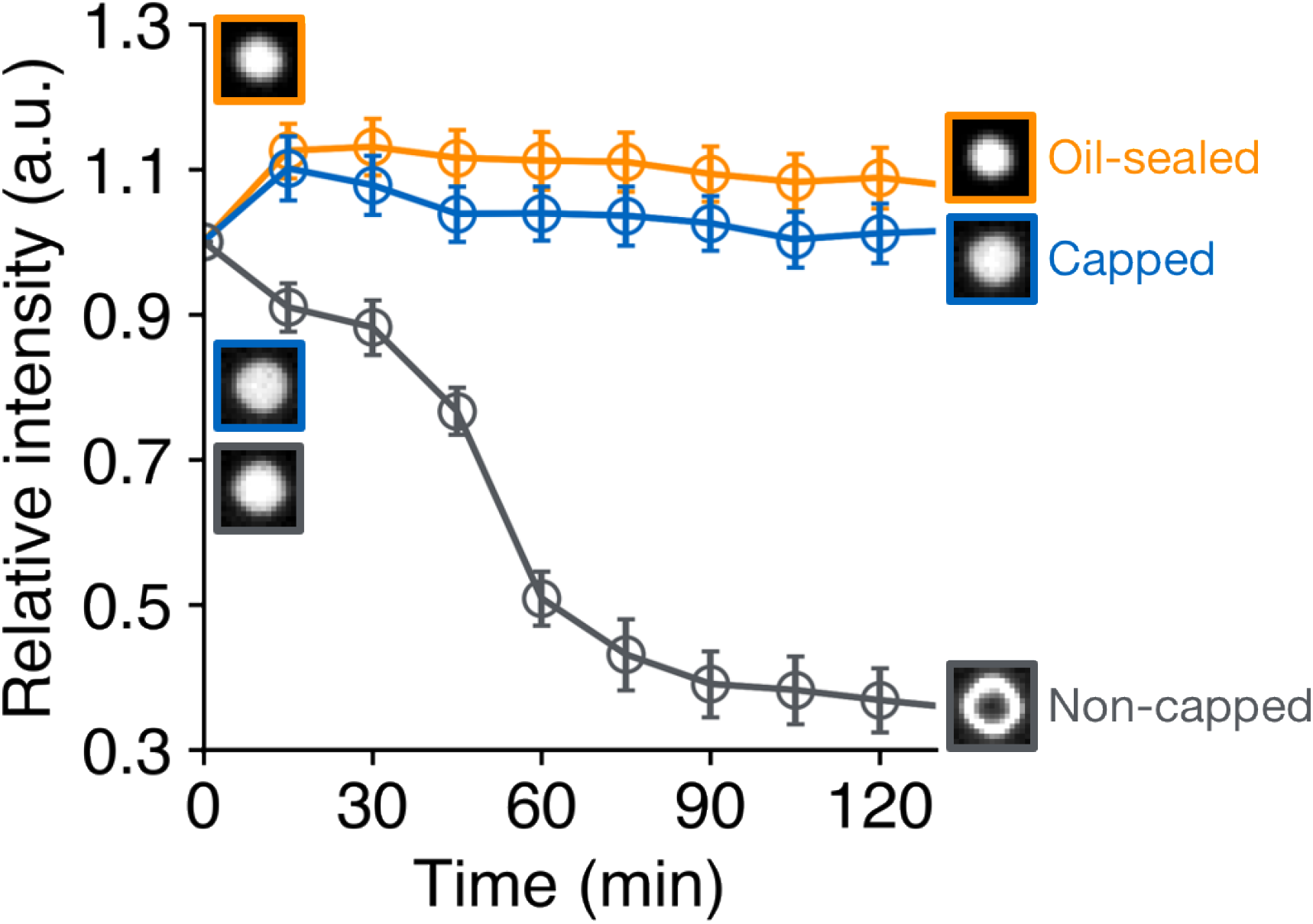
Solution stability in air-sealed reactors. The mean ± SD of the fluorescence intensity of Alexa Fluor 647 at the center of each reactor is plotted. The fluorescence intensity is normalized to that at t = 0 min. Air-sealed reactors with capped flow channel (blue), with non-capped flow channel (gray), and oil-sealed reactors (orange) are shown. *N* = 1053 for each condition.

**Fig. S2.**
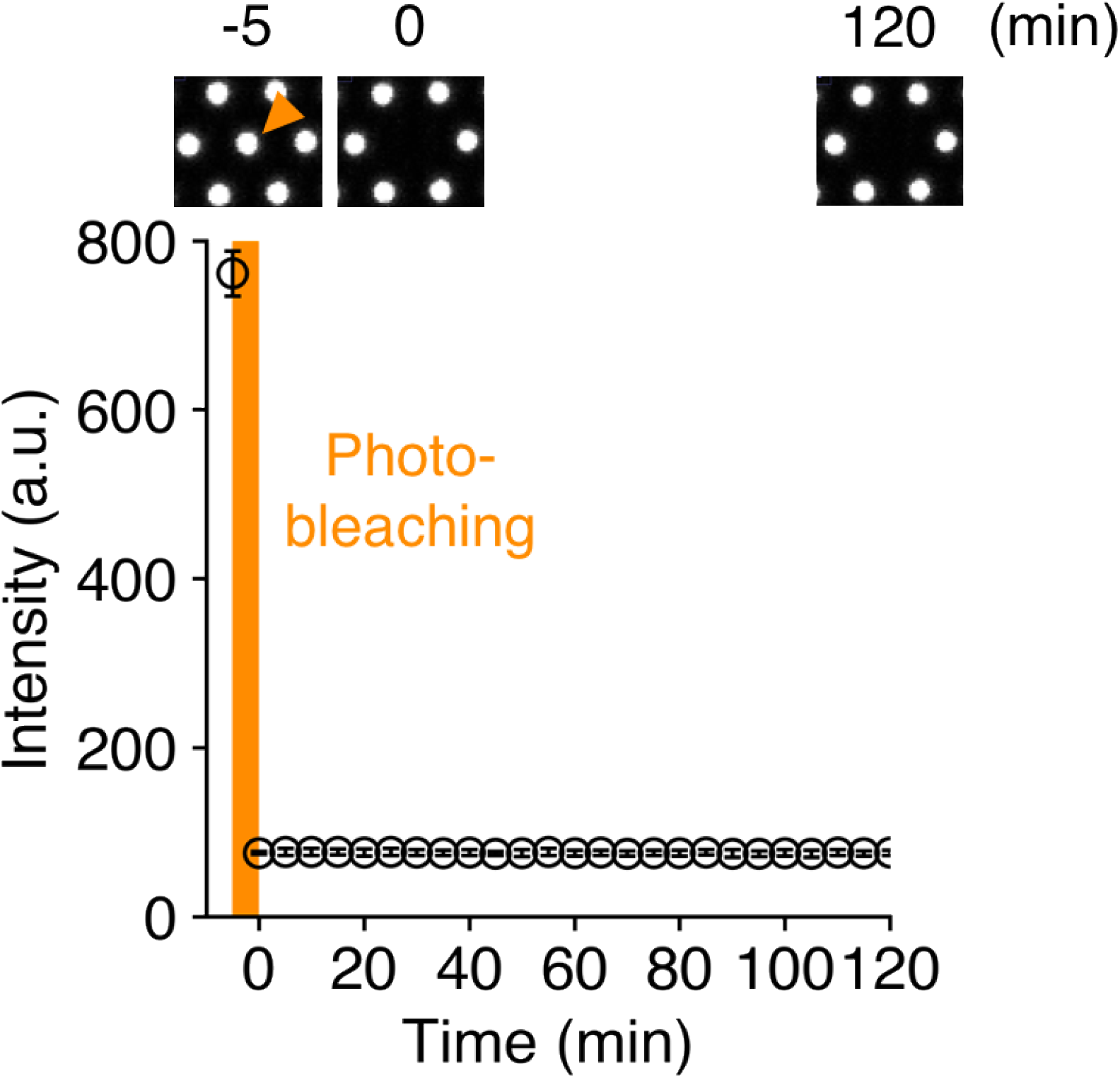
Solution independence in air-sealed reactors. Reactors are photobleached at t = −5 min. The mean ± SD of the fluorescence intensity of Alexa Fluor 647 in photobleached reactors is plotted. *N* = 10. The top panels show fluorescent images of a reactor before (t = −5 min) and after photobleaching (t = 0 and 120 min).

**Fig. S3.**
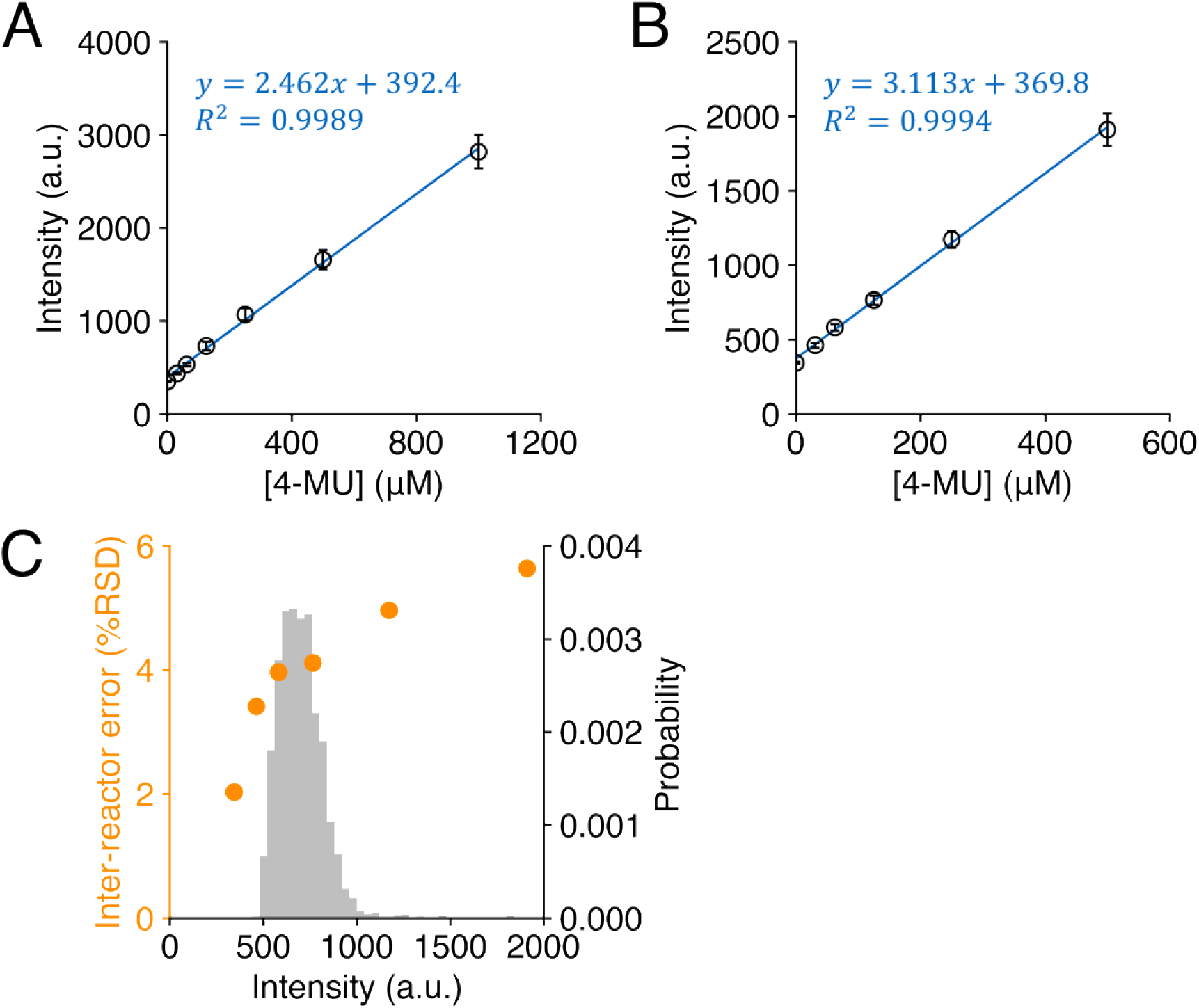
Calibration curve of 4-MU in assay buffer A (A) and assay buffer B (B). The mean ± SD of the fluorescence intensity in the reactors is plotted against the dose of 4-MU. (A) *N* > 190000 and (B) *N* > 220000. (C) Inter-reactor error (corresponding to % RSD of the fluorescence intensity in [B]) is plotted against the fluorescence intensity of 4-MU in a reactor (orange). The gray histogram shows the distribution of fluorescence intensity of 4-MU in reactors at t = 4 min in the NA turnover measurement.

**Fig. S4.**
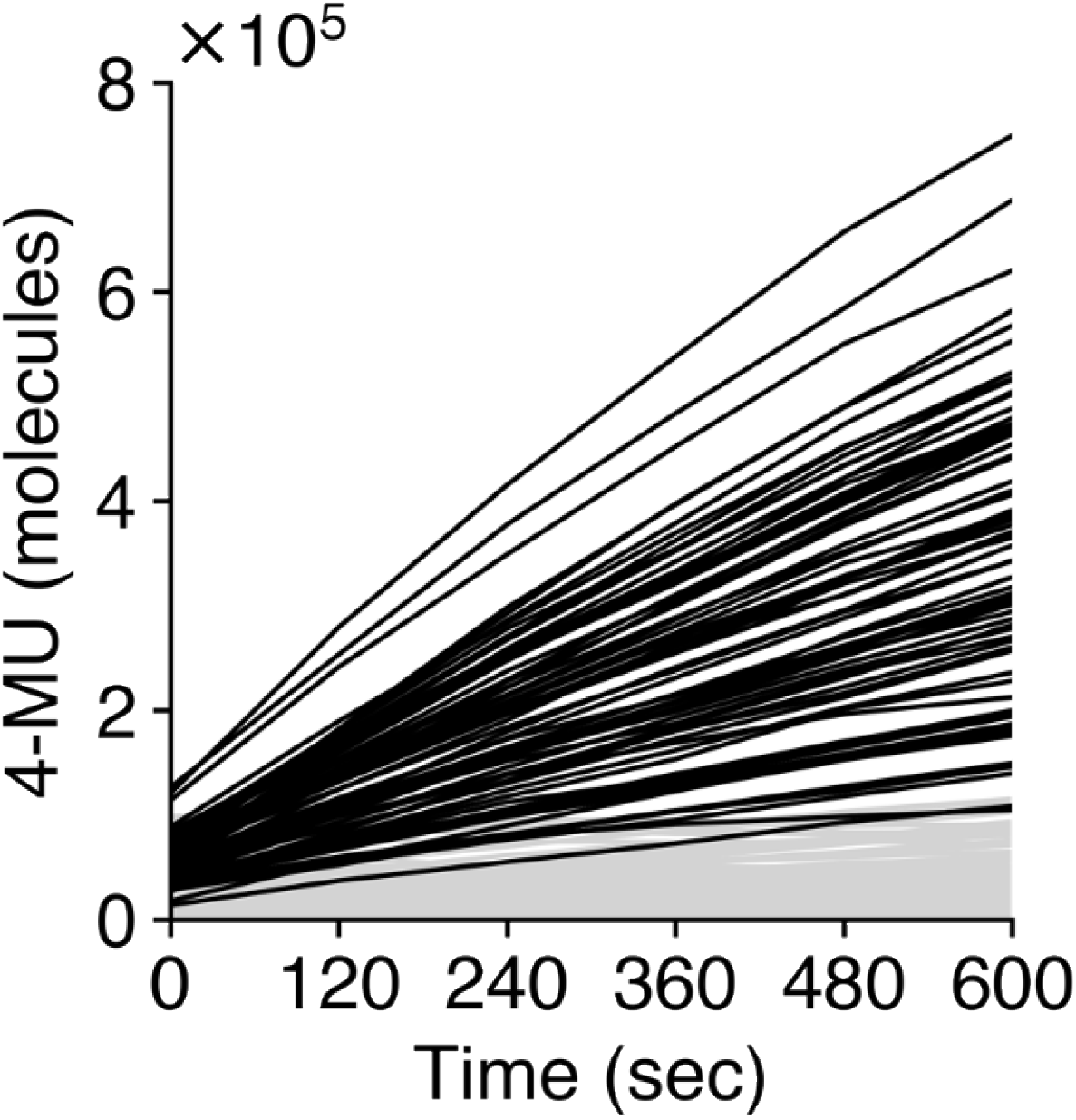
Change in the number of 4-MU molecules in typical reactors over time. Black and gray plots represent IAV particles of which NA turnovers are above and below the threshold, respectively, as defined in Fig. 2C.

**Fig. S5.**
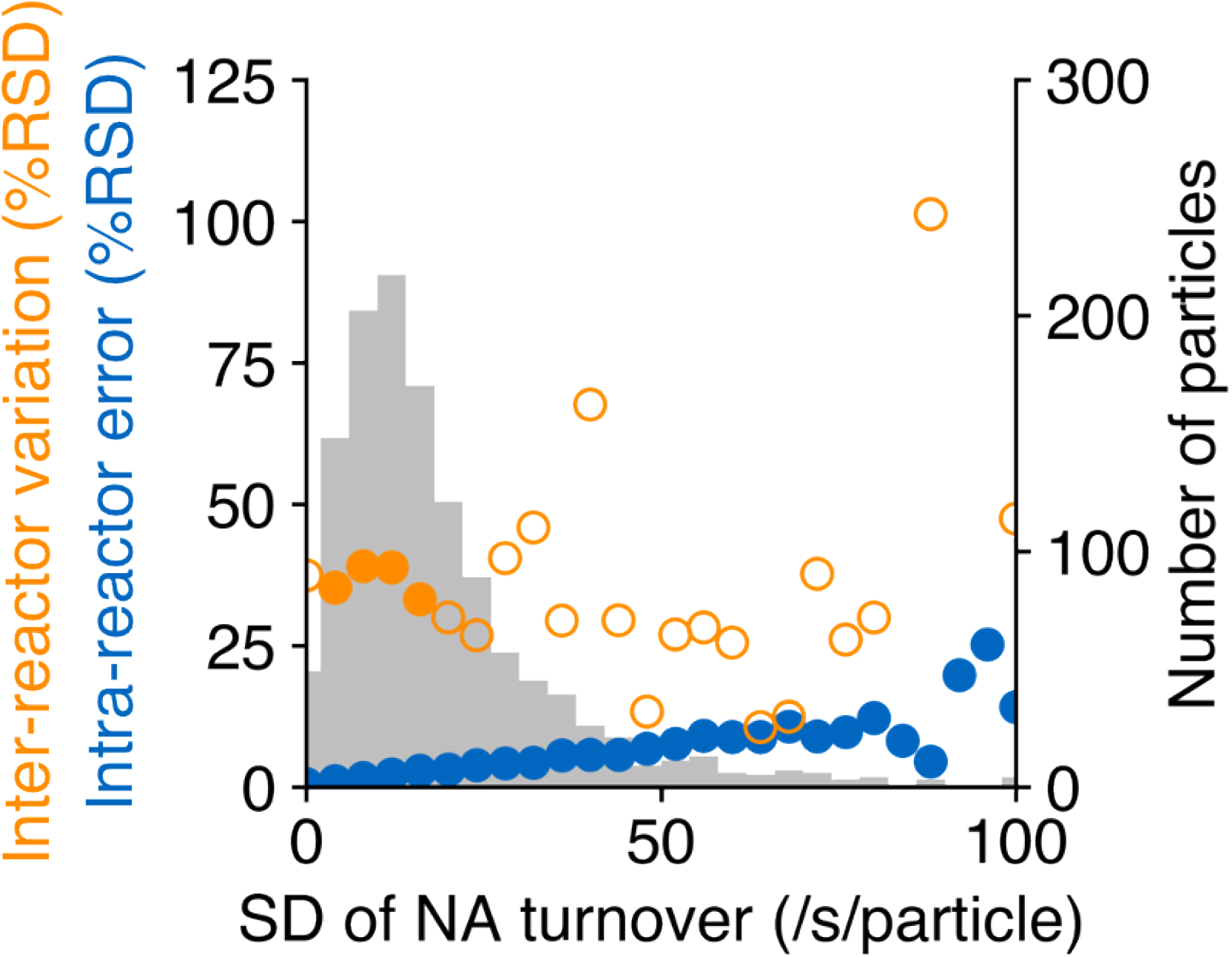
Inter-reactor variation and intra-reactor error in NA turnover measurement estimated by one-way ANOVA. Gray: number of virus particles in each sub-population sliced with the SD of NA turnover measurement. Blue: intra-reactor error of each sub-population. Orange: inter-reactor variation in each sub-population (Sub-populations that have more than 10% of the original population are shown in closed circles).

**Fig. S6.**
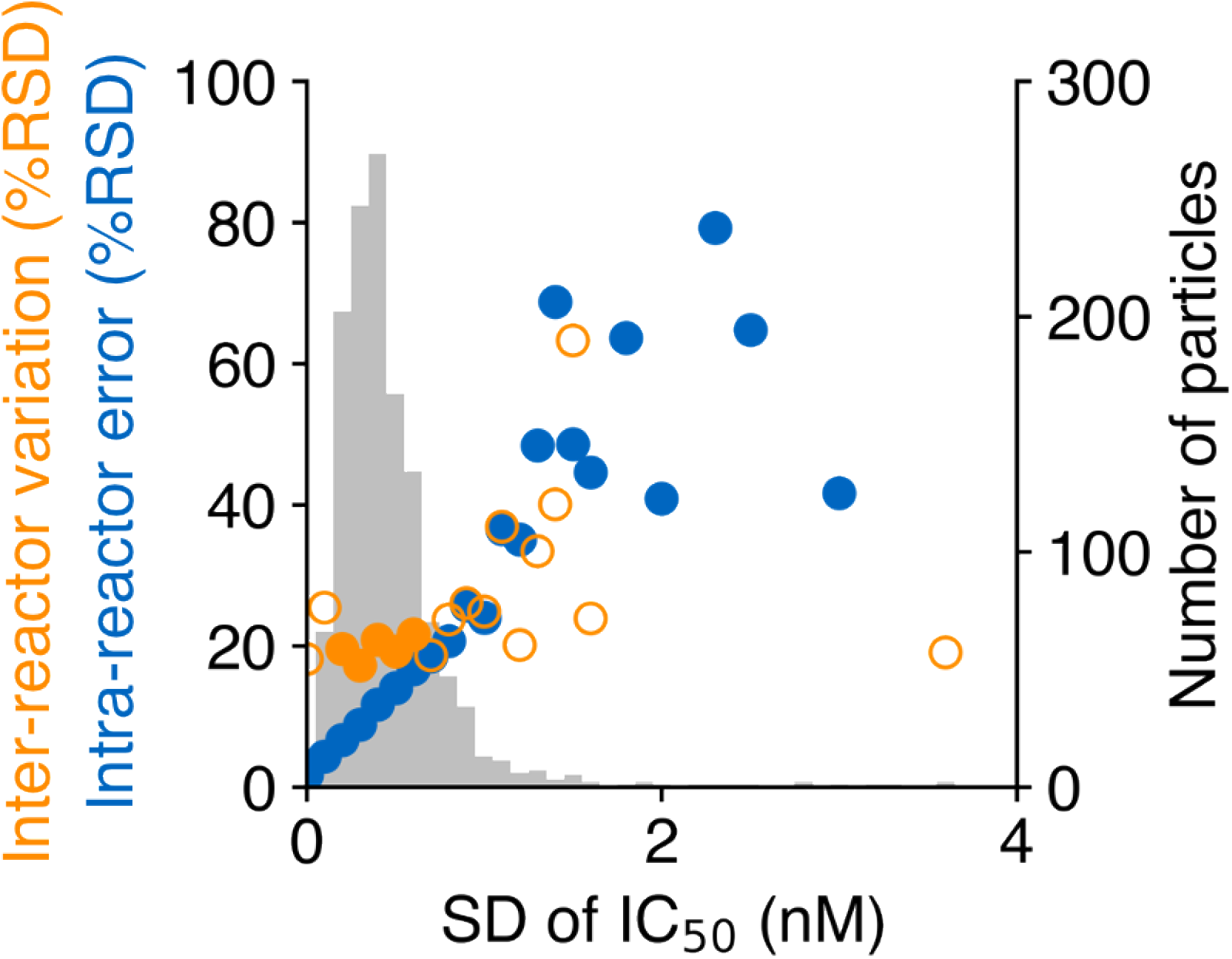
Inter-reactor variation and intra-reactor error in IC_50_ measurement of oseltamivir estimated by one-way ANOVA. Gray: number of virus particles in each sub-population sliced with the SD of the IC_50_ measurement. Blue: intra-reactor error in each sub-population. Orange: inter-reactor variation in each sub-population (Sub-populations that have more than 10% of the original population are shown in closed circles).

**Fig. S7.**
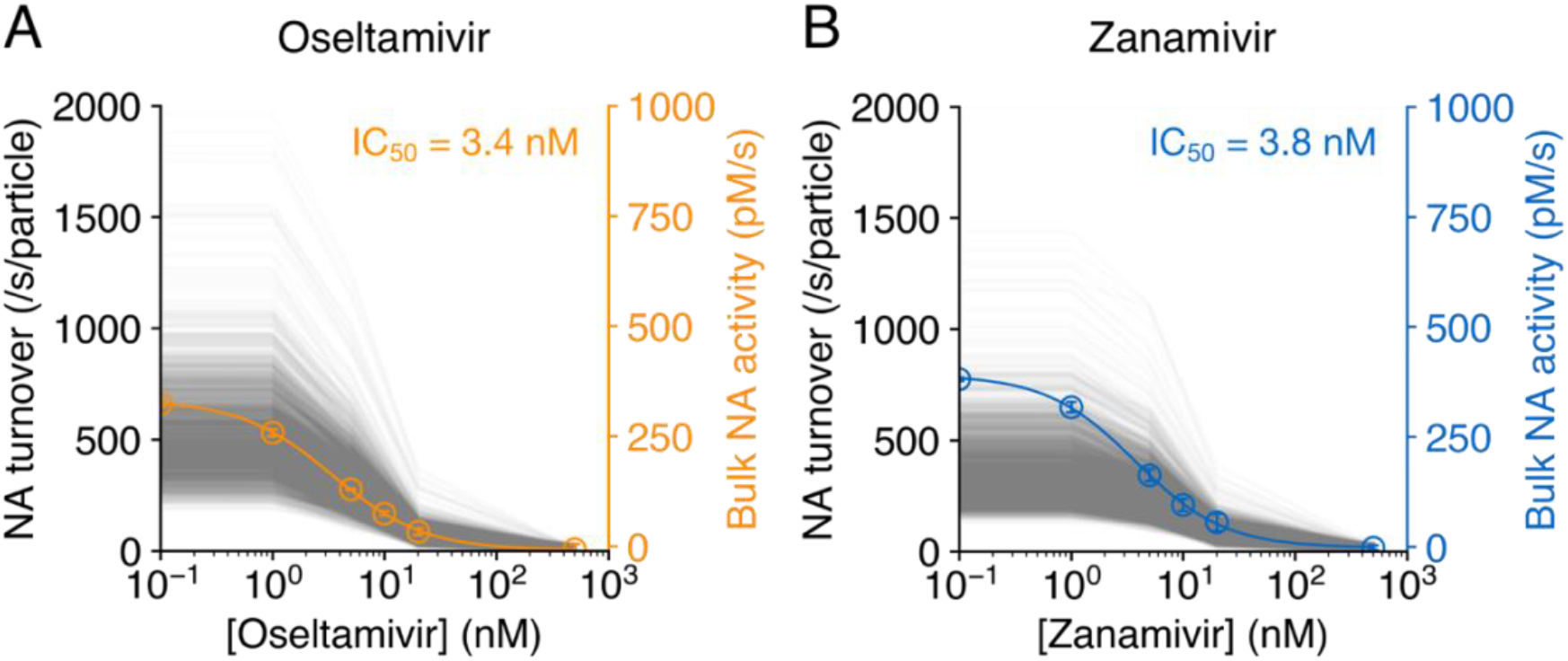
NAI dose-dependency of NA turnover. (A) Oseltamivir or (B) zanamivir dose-dependent decrease in NA turnover is shown for 2286 virus particles. Each gray plot represents a single virus particle. The mean ± SD of NA turnover measured in a conventional bulk scale bioassay is plotted versus (A) oseltamivir or (B) zanamivir concentration and fitted with the Hill equation to calculate the IC_50_. The resulting IC_50_ is shown in each figure.

**Fig. S8.**
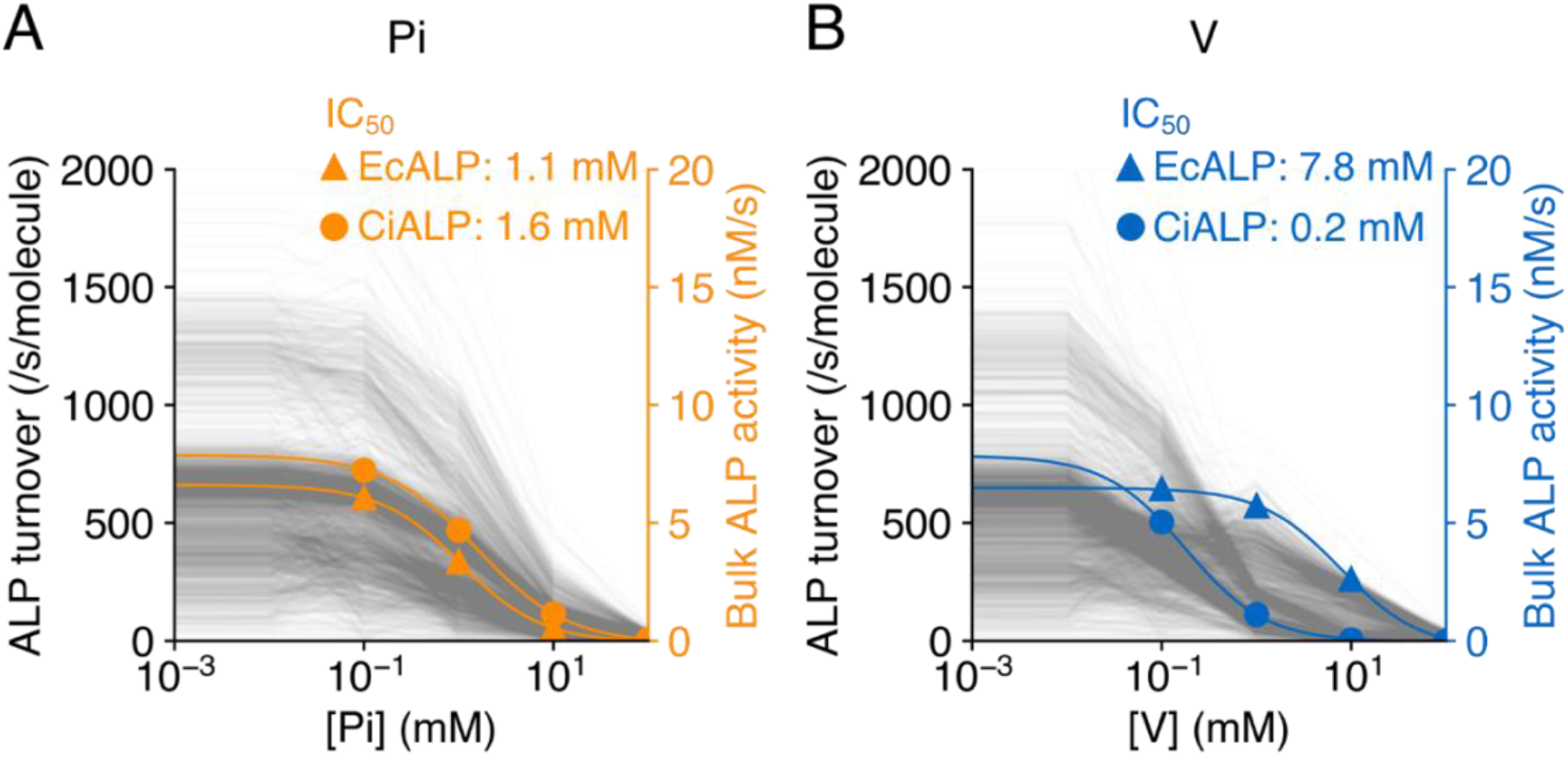
Inhibitor dose-dependency of ALP turnover. (A) Pi or (B) V dose-dependent decrease in ALP turnover is shown for 2200 enzyme molecules of ALP mix. Each gray plot represents a single enzyme molecule. The mean ± SD of ALP turnover measured in a conventional bulk scale bioassay of EcALP or CiALP is plotted versus (A) Pi or (B) V concentration and fitted with the Hill equation to calculate the IC_50_. The resulting IC_50_ is shown in each figure.

**Fig. S9.**
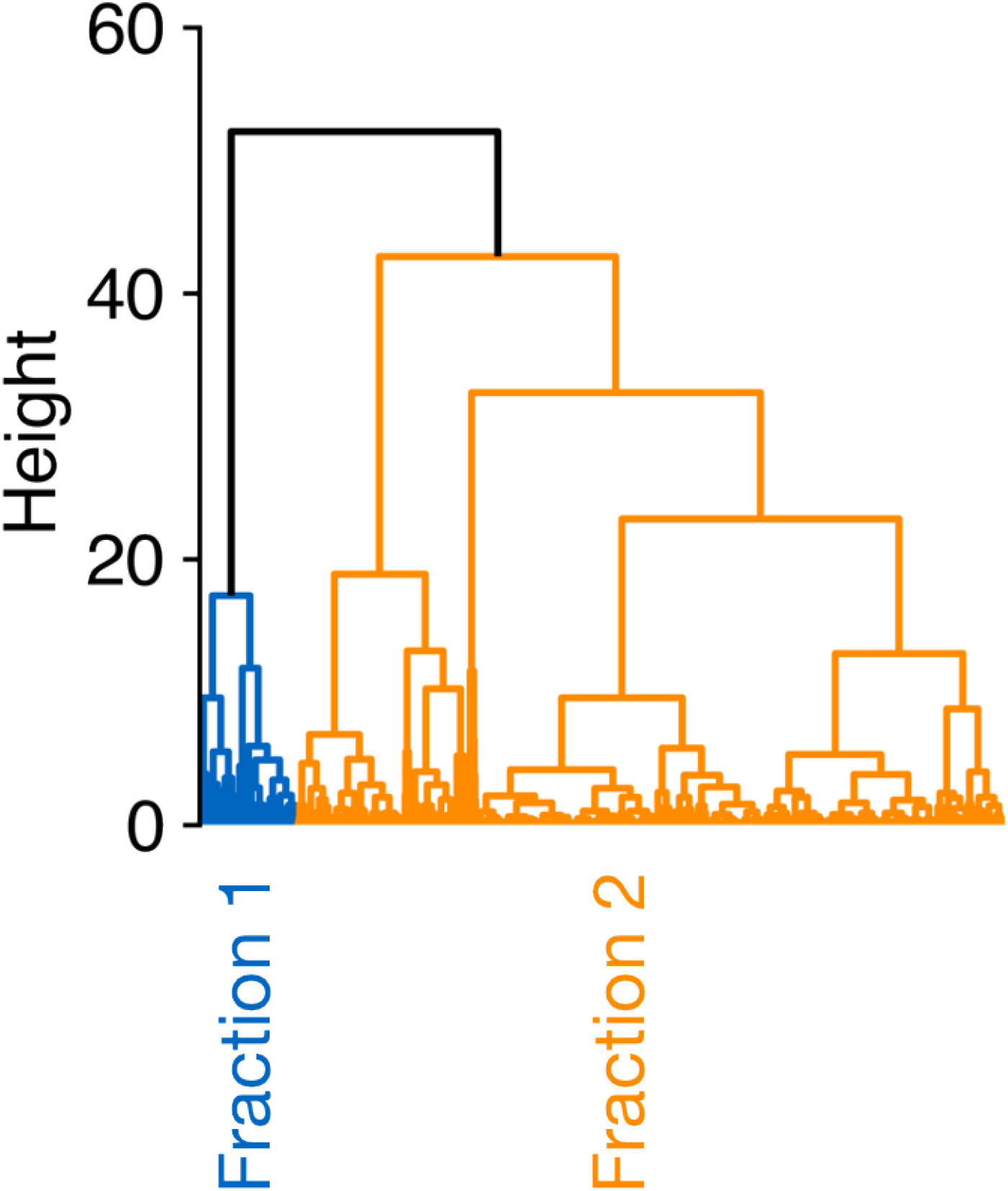
Dendrogram generated by cluster analysis of ALP mix.

